# Direct aliquoting from a single heated well enables fully automated high-throughput thermal proteome profiling

**DOI:** 10.64898/2026.09.15.751689

**Authors:** Mirva Pääkkönen, Iris Purma, Johannes Merilahti, Otto Kauko

## Abstract

Thermal proteome profiling (TPP) measures drug target engagement across the proteome by detecting ligand-induced changes in protein thermal stability, but conventional workflows are laborious and costly, which has limited their use in compound screening. We present DASH-PISA, a single-well thermal fractionation approach in which a lysate is heated through a series of defined temperatures in one well and sampled at each step, and the aliquots are pooled into another well. Because sampling and pooling both take place in plate format, the thermal treatment runs on a standard liquid-handling robot with an integrated thermocycler and a 96-channel pipette. Combined with filter-based separation of soluble protein, data-independent acquisition (DIA), and a spike-in SILAC reference, DASH-PISA forms a fully automated, high-throughput TPP workflow. As a proof of concept, stepwise single-well fractionation reproduced the melting curves of traditional TPP, and the pooled DASH-PISA workflow detected known kinase targets of staurosporine; a spike-in SILAC reference further improved their recovery. By running on commercially available automation, this workflow makes proteome-wide target identification practical at the scale required for compound-library screening.

## Introduction

Most drugs are designed to act on a single protein, yet few are truly selective. Nearly all compounds also engage unintended proteins, and these off-target interactions can be harmful or, occasionally, beneficial. Thalidomide, for example, was prescribed for morning sickness in the 1950s and 1960s and caused severe birth defects through its binding to cereblon.^1,2^ The same interaction underlies its later use as an anticancer agent.^3^ Systematic measurement of the proteins a compound engages, both intended and unintended, is therefore central to modern drug discovery.

Thermal proteome profiling (TPP) measures target engagement directly in cell lysates or intact cells by exploiting protein thermal stability.^4^ Most proteins denature and aggregate when heated, and ligand binding typically stabilises the soluble folded state, shifting the temperature at which a protein becomes insoluble. In a TPP experiment, the sample is heated across a range of temperatures, soluble protein is separated from aggregates, and the soluble fraction is quantified by mass spectrometry (Figure 1A). TPP has proven powerful but also laborious and costly, which has limited its use to focused, hypothesis-driven studies. Proteome integral solubility alteration (PISA) reduces this burden by pooling the temperature fractions of each sample before analysis, which substantially lowers the number of mass spectrometry measurements and removes the need to fit melting curves (Figure 1B).^5^ Replacing ultracentrifugation with filter-assisted separation of soluble protein further allows the assay to run in 96-well plates.^6,7^

**Figure 1.**
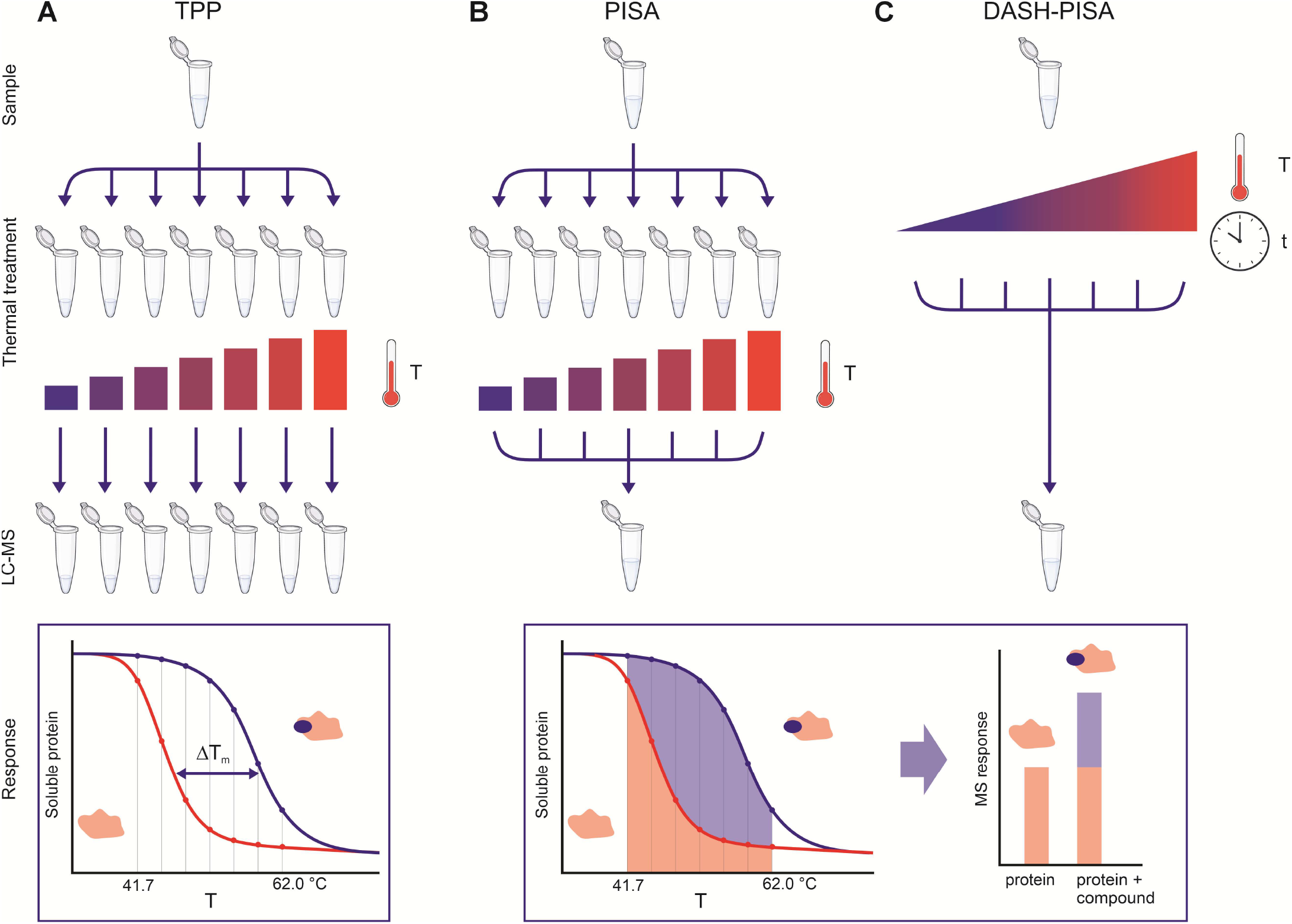
Comparison of thermal-treatment strategies for TPP. (A) Traditional TPP: a sample is divided into separate aliquots, each aliquot is heated to a different temperature, and every aliquot is analysed individually. In traditional TPP, drug-induced ΔT_m_, the shift in melting temperature between the treated and non-treated protein, is determined by sigmoid curve fitting. (B) PISA: aliquots are heated to different temperatures as in traditional TPP, but after heating they are pooled into a single sample that reports the integrated soluble fraction across the temperature range. In PISA melting curve is not required. Instead, the relative quantities of soluble proteins are determined from protein signal abundances in treated and non-treated samples. (C) DASH-PISA: a single well is heated stepwise through a series of defined temperatures, an aliquot is collected at each step, and the aliquots are pooled into a second plate. Colour indicates temperature. Because both sampling and pooling take place in plate format, the DASH-PISA thermal treatment can be fully automated. In DASH-PISA, relative quantities of soluble proteins are determined in the same way as in PISA.

Temperature fraction pooling and plate-based processing reduce the labour of TPP, but quantitation is a separate problem that determines how reliably small stability changes can be detected. Tandem mass tag (TMT) labelling has been the standard quantitation strategy for TPP because multiplexing provides an internal reference that controls for experimental and analytical variation. This is particularly important in TPP because target-induced stability changes are often small and their magnitude does not necessarily reflect binding affinity or biological importance.^4^ However, TMT labelling complicates sample preparation, requires offline fractionation for deep coverage, and is difficult to automate. Data-independent acquisition (DIA) provides a label-free alternative to TMT. It requires simpler sample preparation, is compatible with automation, and, on current instruments, reaches deep proteome coverage from single injections with short gradients. Other recent studies reached the same conclusion and applied DIA to TPP and PISA.^8,9^ DIA’s drawback relative to TMT labelling is the loss of a built-in internal reference. Adding a spike-in reference labelled with heavy amino acids (stable isotope labelling by amino acids in cell culture, SILAC) supplies that control and improves quantitative accuracy,^10^ while retaining the simplicity of DIA. A spike-in reference benefits DIA-based TPP even when unlabelled,^11^ but isotope labelling pairs each reference peptide with its endogenous counterpart, giving a direct per-peptide ratio and thus a more accurate estimate of sample abundance relative to the reference.

Automating the full workflow would open TPP to compound-library screening, but the thermal-treatment step has remained an obstacle because each sample is conventionally split into many separate aliquots and heated to different temperatures. Recent high-throughput PISA workflows have streamlined the sample preparation and DIA analysis that follow thermal treatment,^12^ yet the thermal treatment itself still relies on splitting each sample and heating the aliquots separately. Aliquots can instead be collected from a single reaction as it is heated. In stepwise single-well thermal fractionation, one well is heated stepwise through a series of defined temperatures and an aliquot is collected at each step. Analysing these fractions individually reconstructs a melting curve. Pooling them into a second plate gives a PISA-style measurement, which we call DASH-PISA (Direct Aliquoting from a Single Heated well; Figure 1C). Sampling across a range of temperatures matters because proteins differ widely in the temperature at which they aggregate, and a single temperature reports mainly on those that destabilise near it. A single reaction has been sampled repeatedly before, in one-pot time-induced PISA, where the sample is held at a constant temperature in a heated capillary and fractions are separated by residence time.^13^ That approach is isothermal and depends on custom-built instrumentation. Stepwise fractionation instead raises the temperature between samplings and runs on a standard thermocycler. Because both sampling and pooling take place in plate format, the procedure maps directly onto a robot with an integrated thermocycler and a 96-channel pipette. The complete assay is therefore straightforward to automate. We combine DASH-PISA with filter-based separation, a spike-in SILAC reference added after the thermal treatment, and DIA, giving a fully automated, high-throughput TPP workflow. We show that stepwise single-well fractionation reproduces the melting curves of traditional TPP, that DASH-PISA detects known kinase targets of staurosporine, and that spike-in SILAC improves the recovery of known targets.

## Experimental Section

### Cell lysis and drug treatment

HEK293FT cells (Thermo Fisher Scientific) were cultured in Dulbecco’s modified Eagle’s medium (DMEM, high glucose; D6171, Sigma-Aldrich) supplemented with 10% (v/v) fetal bovine serum, 2 mM L-glutamine, 100 U/ml penicillin, and 100 µg/ml streptomycin. Cultures were maintained at 37 °C in a humidified atmosphere containing 5% CO2 and were passaged before reaching confluence.

For the spike-in SILAC experiment, both light and heavy cells were cultured in DMEM for SILAC (Thermo Fisher Scientific, 88364) supplemented with 10% (v/v) dialysed fetal bovine serum, 2 mM L-glutamine, and either light or heavy L-lysine and L-arginine, respectively (Thermo Fisher Scientific). The heavy amino acids were [13C6, 15N2]-L-lysine (Lys8) and [13C6, 15N4]-L-arginine (Arg10). Cells were cultured in SILAC medium for at least ten population doublings before the experiment. Cells were harvested at approximately 80% confluence by trypsin, pelleted, resuspended in ice-cold phosphate-buffered saline (PBS) supplemented with protease inhibitors (cOmplete, Roche), and distributed into a 96-well PCR plate. Cells were lysed by five cycles of freezing in liquid nitrogen and thawing at 35 °C. Lysates were incubated with 10 µM staurosporine or an equivalent volume of vehicle (DMSO) as a control for 30 min at 37 °C in the 96-well PCR plate.

### Thermal fractionation by direct aliquoting from a single heated well

Stepwise single-well thermal fractionation was performed on an Opentrons Flex robot (Opentrons Labworks). For the melting curve experiment, lysates were heated in the integrated thermocycler module across seven temperature points. Samples were first heated to 41.7 °C and held for 3 min, after which an aliquot of each was transferred to a fresh PCR plate. The same wells were then heated successively to 44.5, 48.0, 51.5, 55.0, 58.5, and 62.0 °C, and an aliquot was collected at each step; each temperature fraction was analysed individually to reconstruct melting curves. For the spike-in SILAC DASH-PISA experiment, light-labelled lysates were fractionated in the same stepwise manner across three temperatures, 52.0, 54.5, and 57.0 °C, and the aliquots were pooled. After pooling, an equal volume of heavy-labelled reference lysate was added to the pooled light-labelled sample and mixed.

### Separation of soluble protein and digestion

Heated fractions from the melting curve experiment and samples from the spike-in SILAC experiment were transferred to a 96-well filter plate (0.45 µm pore size, Millipore) and centrifuged at 1 500 g for 10 min at +4 °C to separate soluble protein from aggregates.^6,7^ The filter plate was washed with 15 µl of ice-cold PBS containing protease inhibitors, and the centrifugation was repeated. Soluble protein was digested by protein aggregation capture, modified from Batth et al. (2019)^14^ and Kverneland et al. (2024)^15^. Briefly, proteins were treated with 1% sodium dodecyl sulphate (SDS), reduced with 5 mM tris(2-carboxyethyl)phosphine (TCEP), and alkylated with 10 mM chloroacetamide (CAA). Proteins were aggregated onto magnetic beads (MagReSyn Hydroxyl beads, ReSyn Biosciences) by adding acetonitrile, and the samples were processed on an Opentrons Flex robot (Opentrons Labworks). Proteins were digested overnight with Mass Spec Grade Trypsin/Lys-C Mix (Promega). Digested peptides were loaded onto Evotips (Evosep, Odense, Denmark) using an Opentrons Flex robot.

### LC-MS/MS

LC-ESI-MS/MS analysis was performed on an Evosep One HPLC system (Evosep, Odense, Denmark) coupled to an Orbitrap Astral or Orbitrap Astral Zoom mass spectrometer (Thermo Scientific, Bremen, Germany) equipped with a nano-electrospray ionisation source. Peptides were separated in-line on an Aurora Rapid C18 UHPLC column (8 cm × 150 µm, IonOpticks, Collingwood, Australia). The mobile phase consisted of water with 0.1% formic acid (solvent A) and 0.1% formic acid in 99.9% acetonitrile (v/v) (solvent B). Samples were analysed with a 100 samples-per-day method using data-independent acquisition (DIA). Data were acquired automatically with Thermo Xcalibur (Thermo Scientific) software version 4.7 (Orbitrap Astral) or 4.8 (Orbitrap Astral Zoom).

On the Orbitrap Astral and Orbitrap Astral Zoom, MS1 spectra were collected in the Orbitrap mass analyser every 0.6 s, and each duty cycle contained one full scan over 380–980 m/z at a resolution of 240 000, with the normalised full-MS AGC target set to 3e6 (Orbitrap Astral) or 5e6 (Orbitrap Astral Zoom). For the Orbitrap Astral, MS2 ions were scanned over *m/z* 380.428–980 using 167 DIA MS/MS scans with variable-width isolation windows. For the Orbitrap Astral Zoom, MS2 ions were scanned over *m/z* 380–980 with 2.5 Da isolation windows. For both instruments, MS2 fragmentation used a normalised collision energy of 25%, and fragment scans were recorded with a maximum fill time of 2.5 ms (Orbitrap Astral) or 2.0 ms (Orbitrap Astral Zoom). The normalised AGC target was set to 10 000 (Orbitrap Astral) or 50 000 (Orbitrap Astral Zoom).

### Data analysis

Raw data were processed with DIA-NN 2.5.1 against the *Homo sapiens* database (SwissProt release 2026_01). Trypsin/P was set as the enzyme with one allowed missed cleavage. Carbamidomethylation of cysteine was set as a fixed modification, and oxidation of methionine and excision of N-terminal methionine were set as variable modifications. For the spike-in SILAC experiment, heavy and light channels were quantified separately for ratio-based normalisation. The light channel was set to lysine 0 and arginine 0, and the heavy channel was set to lysine 8.0142 and arginine 10.0083. Peptide length was restricted to 7–35 residues and precursor charge to 2–4. Results were filtered at a false discovery rate (FDR) of 1%.

Melting curve analysis was performed using the DIA-NN protein-group report. Protein groups were mapped to gene symbols, and abundances were log2-transformed. Proteins annotated as non-melters are expected to remain soluble across the temperature range and were therefore used as a normalisation reference. Abundances were normalised to a non-melter reference derived from the Meltome Atlas^16^. The reference comprised proteins annotated as non-melters in at least 20 human datasets and quantified in our data. For each condition and temperature, the median log2 abundance across these proteins was calculated, and all abundances were centred by subtracting it. The normalised abundance was expressed as the soluble fraction relative to the lowest measured temperature, so that the curves started near 1. Plotted points represent the median soluble fraction across replicate measurements. Curves were fitted per protein and condition by nonlinear least squares (nlsLM, minpack.lm) to a three-parameter sigmoid with the upper plateau fixed at 1, y(T) = bottom + (1 − bottom) / (1 + exp((T − Tm) / scal)), where bottom is the fitted lower plateau and scal determines the steepness of the transition. The melting temperature (Tm) was taken as the fitted midpoint.

Kinases were defined as the human proteins annotated with the UniProt keyword Kinase (KW-0418, organism 9606). Kinases highlighted in the figures and reported as targets were manually reviewed to retain only protein kinases, the expected target class of staurosporine. Staurosporine-induced changes in soluble abundance were assessed using two-tailed t-test, and p-values were adjusted with the Benjamini-Hochberg procedure. Protein kinases were reported as targets at q < 0.01 and q < 0.05.

## Results and Discussion

The central obstacle to automating TPP is the thermal-treatment step. In traditional TPP, each sample is divided into aliquots, each aliquot is heated to a different temperature, and every aliquot is analysed separately, which multiplies the number of samples and mass spectrometry runs (Figure 1A). PISA removes the separate analyses by pooling the aliquots of a sample after heating, so that a single pooled sample reports the integrated soluble fraction across the temperature range (Figure 1B).^5^ Pooling reduces the number of measurements, but the aliquots are still prepared and heated individually, which is awkward to automate in plate format. DASH-PISA performs this step in a single well (Figure 1C). The sample is heated stepwise through a series of defined temperatures in one well, an aliquot is withdrawn at each temperature, and the aliquots are pooled into a second plate. Because the aliquots are taken from one plate and deposited into another, the entire thermal-treatment step can be carried out by a liquid-handling robot with an integrated thermocycler and a multichannel pipette.

We implemented DASH-PISA as part of a fully automated, plate-based workflow (Figure 2). Cell lysates in a 96-well plate were treated with compounds of interest, and the thermal treatment and pooling were performed on an Opentrons Flex robot with an integrated thermocycler and a 96-channel pipette. A heavy-labelled spike-in SILAC reference was added to each pooled sample after the thermal treatment to serve as an internal quantitative standard. Soluble protein was separated from aggregates on a 96-well filter plate,^6,7^ and the recovered protein was digested by protein aggregation capture^14,15^ on an Opentrons Flex robot. Peptides were loaded into Evotips and analysed by DIA on an Evosep One coupled to an Orbitrap Astral or an Orbitrap Astral Zoom mass spectrometer using a 100 samples-per-day method, and the data were processed with DIA-NN. Every liquid-handling step from lysate to Evotip was performed on standard, commercially available automation in 96-well format, which is what makes the assay practical for screening.

**Figure 2.**
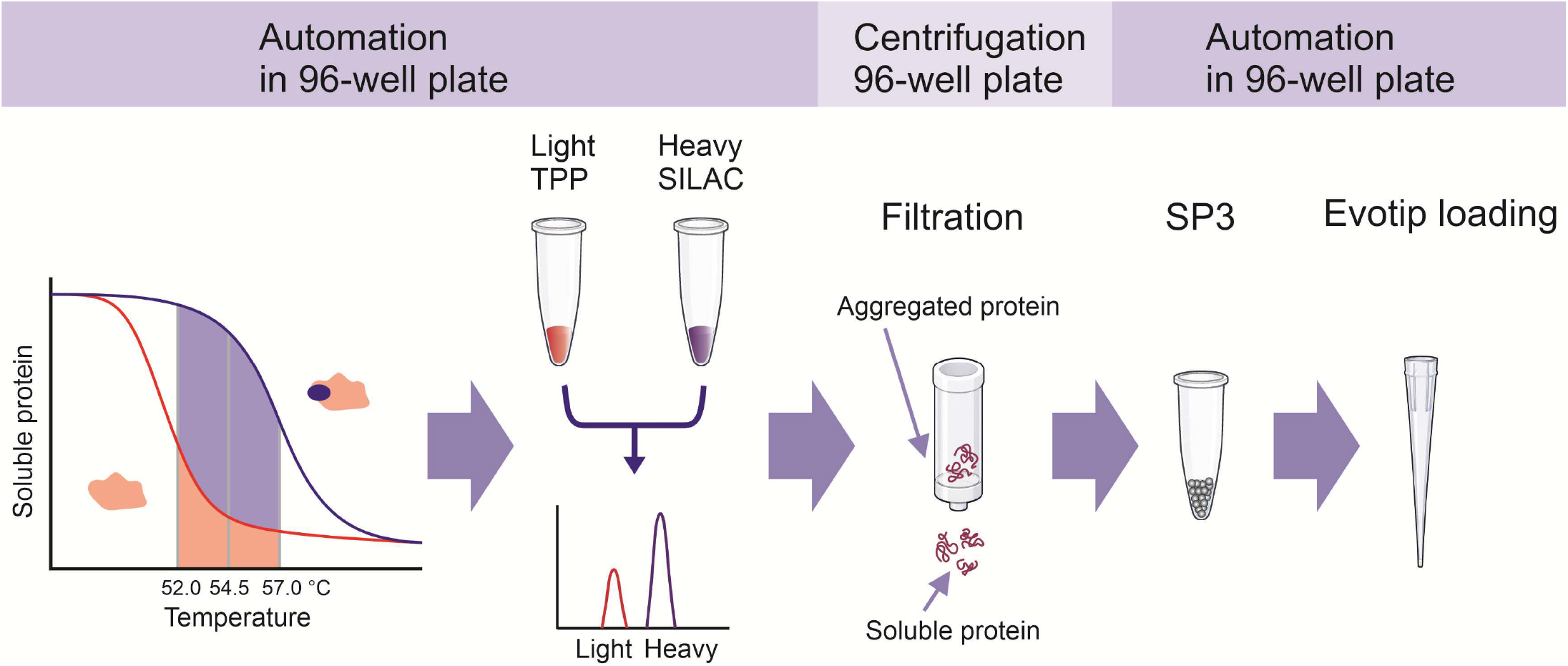
The automated high-throughput TPP workflow. Cell lysates in 96-well plates undergo automated fractionation and pooling (DASH-PISA) on a liquid-handling robot equipped with an integrated thermocycler and a 96-channel pipette. A heavy-labelled spike-in SILAC reference is added to treated samples after the thermal treatment. Soluble protein is separated on a 96-well filter plate, digested by protein aggregation capture on a liquid-handling robot, and loaded into Evotips. Peptides are analysed by data-independent acquisition on an Evosep HPLC system coupled to a fast high-end mass spectrometer. Every liquid-handling step is performed in 96-well format and using standard, commercially available automation.

To test whether repeated sampling of a single well heated in a stepwise manner reproduces conventional thermal behaviour, we compared stepwise single-well fractionation with a traditional TPP experiment on staurosporine-treated lysates, analysing each temperature fraction individually (Figure 3). We normalised each temperature series to proteins annotated as non-melters in the Meltome Atlas^16^, which remain soluble across the heating range and therefore provide a stable reference. Both formats produced sigmoidal melting curves and comparable apparent melting temperatures for representative proteins, including the staurosporine-target kinases ADK, CDK2,^17^ CDK5, CSK,^18^ and GSK3A. Staurosporine stabilised its targets in both assays, shifting the curves to higher temperatures (Figure 3). The stabilisation of CDK2 was particularly pronounced, with an apparent melting temperature shift of about 10 °C. The shifts measured by stepwise single-well fractionation (Figure 3A) agreed with those from traditional TPP (Figure 3B). Both methods gave similar melting temperatures (mean difference of 0.52 °C; two-tailed t-test, p = 0.15, not significant) and staurosporine-induced thermal shifts (mean difference of 0.47 °C; two-tailed t-test, p = 0.45, not significant) across the proteins shown in Figure 3. The close agreement indicates that the cumulative thermal exposure inherent to stepwise sampling does not materially distort apparent thermal stability, and that stepwise single-well fractionation reports comparable target engagement profiles for the proteins examined as the established method.

**Figure 3.**
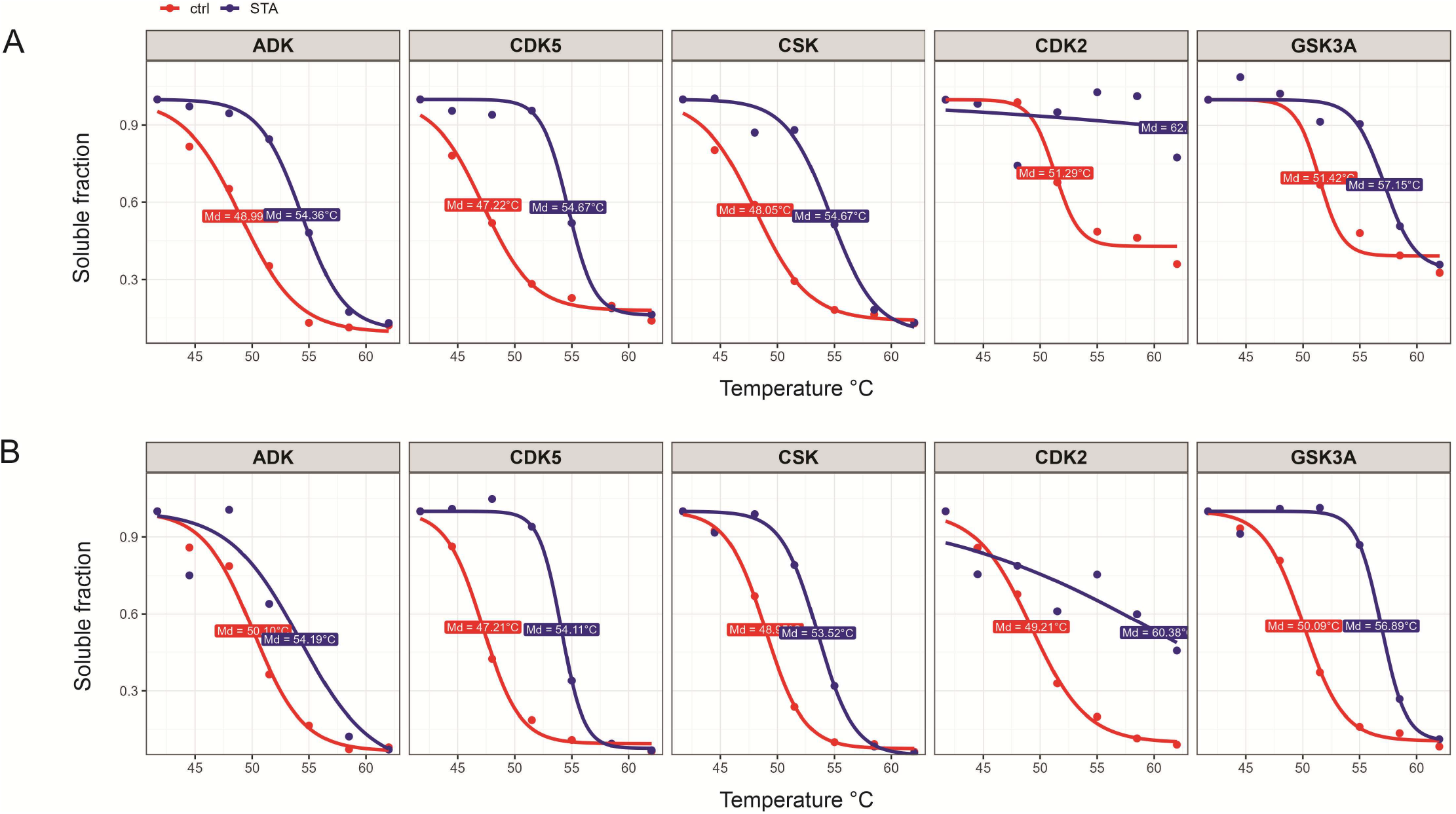
Direct aliquoting from a single heated well reproduces traditional TPP melting curves and detects staurosporine-induced stabilisation. The soluble fraction (relative to lowest T; see methods for details) as a function of temperature for representative proteins in vehicle control (ctrl, red) and staurosporine-treated (STA, blue) samples. (A) Direct aliquoting from a single-well fractionation. (B) Traditional TPP. Points show the median soluble fraction, and curves show the fitted three-parameter sigmoid; the fitted melting temperature (Tm, the sigmoid midpoint, labelled Md) is indicated for each condition. Staurosporine stabilises its kinase targets, for example, CDK2, shifting the curves to higher temperatures in both formats. Data are from six and four replicates for DASH-PISA and traditional TPP, respectively.

Detecting drug targets by TPP requires resolving small changes in soluble abundance against biological and analytical noise. To quantify the benefit of the spike-in SILAC reference, we analysed the same staurosporine dataset with and without SILAC normalisation (Figure 4). Using the light channel alone with median normalisation, 7 protein kinases were identified as significant targets at a Benjamini-Hochberg-adjusted p-value (q-value) below 0.01, and 13 at q < 0.05 (Figure 4A). Normalising each protein to its heavy spike-in reference increased the number of confidently identified protein kinase targets to 14 at q < 0.01 and 16 at q < 0.05 (Figure 4B). Because the reference is added after the thermal treatment, it corrects for variation introduced during protein aggregation capture, digestion, and LC-MS analysis, but not for the thermal treatment itself. The gain therefore reflects improved analytical precision, which is needed to detect the small effect sizes typical of thermal-shift measurements.

**Figure 4.**
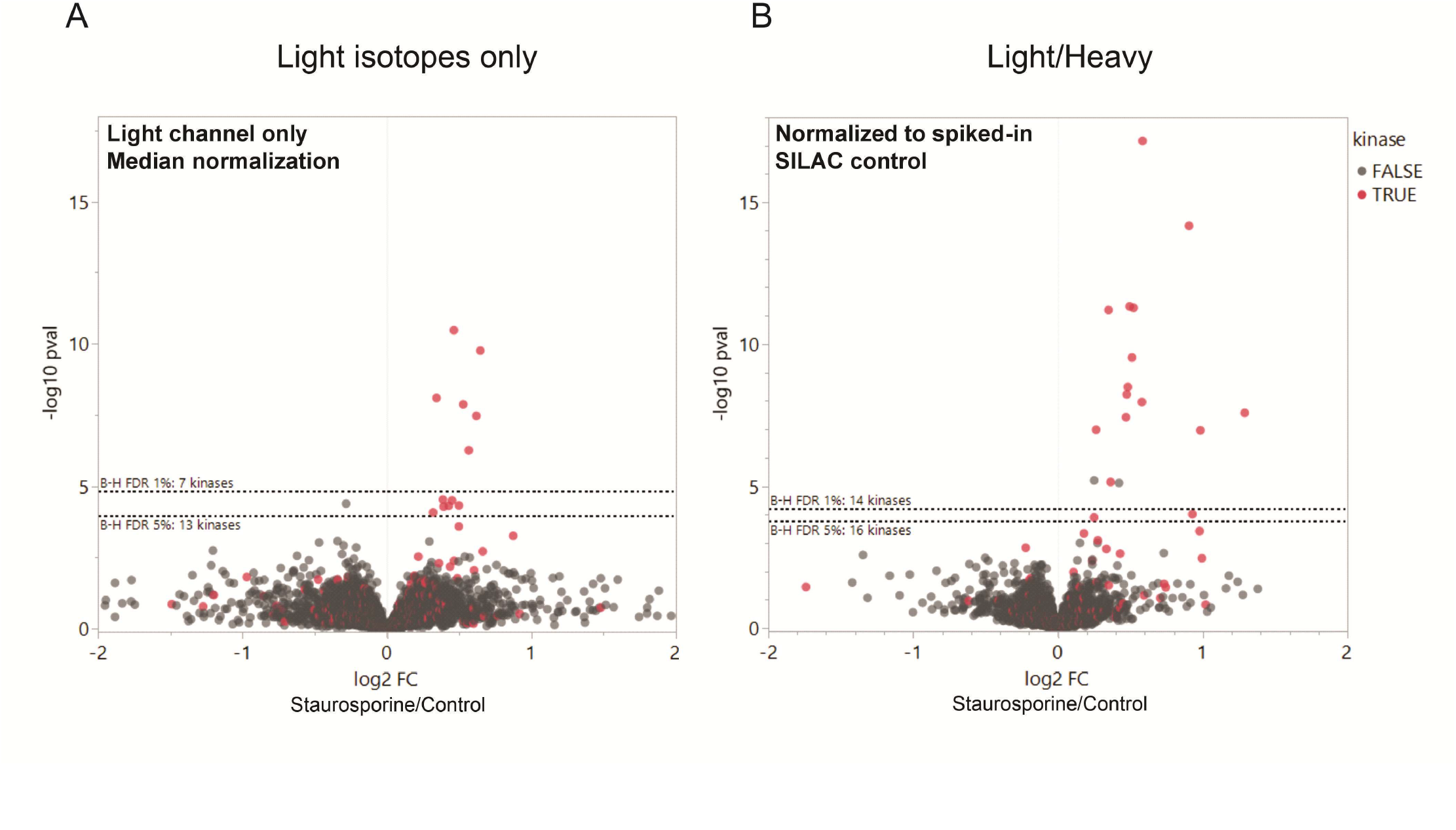
Spike-in SILAC normalisation improves target identification. Volcano plots of the staurosporine versus control comparison, showing the log2 fold change in soluble abundance against statistical significance (-log10 p-value). Protein kinases are shown in red and all other proteins in grey; dotted lines mark Benjamini-Hochberg significance thresholds (q-value). (A) Light channel only with median normalisation: 7 protein kinases at q < 0.01 and 13 at q < 0.05. (B) The same data normalised to the heavy spike-in SILAC reference: 14 protein kinases at q < 0.01 and 16 at q < 0.05. Spike-in SILAC increased the number of confidently identified protein kinase targets (from 7 to 14 at q < 0.01). Two-tailed t-test was used.

Full automation of TPP is enabled here by combining several advances, each addressing a different part of the workflow. A narrow, optimised range of heating temperatures keeps the assay sensitive with only a few temperature points,^6,19^ filter-plate separation replaces ultracentrifugation and moves the assay into 96-well format,^6,7^ and DIA with a spike-in SILAC reference provides accurate, internally controlled quantitation without TMT labelling.^10^ Individually, these advances reduce cost and effort, but the thermal-treatment step still required each sample to be split and heated separately, which prevented full automation of the workflow. DASH-PISA removes that barrier by performing the whole thermal treatment in a single well, and integrating it with the other advances yields a fully automated, plate-based workflow that runs on standard laboratory robotics. The resulting throughput makes TPP practical for compound-library screening and opens the method to questions that were previously out of reach, such as profiling large or unconventional compound collections. The present study is a proof of concept based on the promiscuous inhibitor staurosporine and six replicates; broader validation with more selective compounds, and a controlled comparison of DASH-PISA against static-temperature PISA, in which the sample is held at a constant temperature, will further define the accuracy and limits of the approach.

## Conclusions

We have developed a high-throughput, fully automated thermal proteome profiling workflow built around DASH-PISA, a stepwise single-well thermal fractionation step that removes the last manual barrier to running TPP on standard laboratory robotics. The method reproduces conventional melting curves, identifies known drug targets, and, through spike-in SILAC normalisation, resolves the small thermal shifts relevant to target identification with greater confidence. Because it relies on commercially available instrumentation and DIA-based quantitation, it brings high-throughput TPP within reach of typical proteomics laboratories and makes it feasible to profile large compound libraries. This enables data-driven research questions that were previously impractical, such as characterising the cellular targets of underexplored molecule classes and environmental chemicals.

## Author Information

### Author Contributions

M.P.: investigation, methodology, formal analysis, visualisation, writing - original draft. I.P.: investigation, formal analysis, visualisation, writing - review and editing. J.M.: methodology, writing - review and editing. O.K.: conceptualisation, formal analysis, supervision, writing - original draft. All authors have approved the final version of the manuscript.

### Notes

The authors declare no competing financial interests.

## Acknowledgements

Mass spectrometry analyses were performed at the Turku Proteomics Facility supported by Biocenter Finland.

